# The development of methodology and techniques for crop disease identification

**DOI:** 10.1101/702621

**Authors:** N. S. Tiwari, J. W. Richmond

## Abstract

In India, an estimated 15-25% of potential crop production is lost due pest and diseases (Roy and Bezbaruah, 2002). The country needs not only to raise production but also ensure food security for its growing consumption needs while curbing excessive pesticide usage. Detection of pests and diseases at an early stage plays a significant role in addressing the above-mentioned concerns and image classification offers a cost-effective and scalable solution to the disease detection problem (A. Ramcharan *et al.* 2017). Here, the principles of transfer learning are implemented with pretrained model – Resnet34 (K. He *et al.* 2015), and test its effectiveness in image classification using a dataset of tea leaves. The novelty of this work is that the images used are not curated, individual leaves with controlled backgrounds but of plants in-situ. The effect of the level of zoom and background is examined and class activation maps are used to validate that the basis of classification is indeed the disease and not an artificial bias from factors such as background, lighting etc.

## Introduction

Protecting crop yields from the devastation caused by pest and disease infestations is a very important concern for the food security of humans. With farmland coming under increased pressure, it is becoming vital to detect the onset of diseases and pests at an early stage to reduce the economic loss caused due to the reduction in crop yields. Tackling pests and diseases at an early stage also helps in reducing the amount of pesticides used to bypass all the detrimental effects that overuse of pesticides has for the plant, the soil and future crop yields.

The intention of this work is to develop an analysis procedure to enable farmers to accurately identify different pests and diseases using images taken from smartphone cameras. It was anticipated that this could be done using deep neural networks to carry out classification based on images of crop pest/disease symptoms. In this scenario the neural network would be trained using a dataset of images collected from actual incidents of the diseases (Alfarisy et al. 2018; Kamilaris and Prenafeta-Boldú 2018). Once the model had been trained, it could then be deployed either centrally on a cloud server or locally via an app on a smartphone (if sufficient processing power is available). This paper summarises the work carried out to date and shows that neural networks can indeed provide a very powerful and accurate means of identifying crop disease based on symptoms.

Over recent years deep learning using various architectures has been shown to provide state of the art image classification and object recognition capability (Zhao et al. 2018). Much of this development has been carried out using datasets such as Imagenet (Krizhevsky *et al.* 2012), CocoNet (Lin *et al*. 2014), etc. These datasets, whilst being trained on vast numbers of images and object classes are of limited relevance to many real-world problems since the domain of the classes do not pertain to the specific application. Unfortunately, given the complexity and very large number of parameters of modern image classifiers such as Resnet (Kaiming He *et al.* 2015), DenseNet (G. Huang *et al.* 2016), ResNeXt (S. Xie *et al.* 2017), etc, it is necessary to have hundreds of thousands, even millions of labelled images to train the models. Such large numbers of images are rarely available in real world applications. This issue is overcome by using a process called transfer learning to re-train a neural network that has been trained on one of these large datasets and make it specific and applicable to a new problem domain (S. Ruder, online).

There is a significant body of work where Artificial Neural Networks, Support Vector Machines (SVM) and fuzzy logic have been applied to the domain of crop disease detection using an image dataset by a number of authors (Kaur and Singla 2016; Abdullahi *et al*. 2017). Mohanty et al. used the Plant Village dataset, a public domain image dataset consolidated in a controlled setting, to train deep convolutional neural networks and the performance different network architectures were compared in different experiments based on their F1 scores (Mohanty *et al*. 2016). CaffeNet has been used as a starting point to develop an image classification model on a specially curated dataset of leaves spanning 13 classes (Sladojevic *et al*. 2016; Alfarisy *et al*. 2018). Transfer Learning has also been applied to the purpose of image classification in the crop disease detection domain using field level images of cassava plants. The techniques as applied in the works mentioned above, coupled with remote sensing, IoT, cloud computing and big data analysis can go a long way to address a variety of challenges facing precision agriculture (Kamilaris and Prenafeta-Boldú 2018).

Whilst deep learning has been shown to be a very powerful tool there are known problems that mean that good results can be generated on specific datasets, however, these sometimes do not generalise well and give poor results in production. There are many reasons for this behaviour, the most important of which is the lack of exposure during training of the model to scenarios that the application will encounter in the field. The imbalance in the number of images belonging to different classes in the training dataset, nature of images including attributes like colour, zoom level, angle of image capture, texture among others also introduces bias in the model. The nature of the dataset can also result in overfitting in the model which will be explored in detail when reviewing results of the analyses.

The focus of this paper is to explain the development and validation of a methodology for data capture and processing to avoid the above problems and the initial application of deep learning models for this specific application. The work presented provides a strong foundation for future development of robust and scalable crop disease diagnostic tools.

Since the ultimate intention of this work is to help farmers accurately identify different pests and diseases using images taken from smartphone cameras, the images in the training dataset have been taken in-situ. This is a very different approach than those adopted in previous works such as Mohanty (Mohanty *et al.* 2016), and implies that each image is likely to have a complex set of attributes, background noise, contain multiple leaves and/or plants and more than one type of pest or disease. It is also likely that the farmers will take images under varied lighting conditions, zoom levels and from different angles. Hence, the initial focus was to examine what sort of images would be effective in training the model to make predictions with high accuracy levels. The results demonstrated in this work will be used to establish a baseline for the methodology of dataset collection and its importance in the development of a robust automated disease recognition system.

The experimentation elaborated in this paper was done on the leaves of different sections and growing conditions in a tea plantation. This was done with the support of Amalgamated Plantations Private Limited (APPL is an associate company of Tata Global Beverages) and Mr. Kailyanjeet Borah, R&D manager who provided essential assistance during the data collection exercise, which was carried out in Teok Tea Estate in the Jorhat city of the state of Assam, India.

## Materials and Methods

Targeting the different kinds of pest & disease symptoms in a single plant type, the Tea Crop facilitated consistency in the physical attributes and features of the leaves and provided uniformity in model testing. The pests/ diseases of interest were:

i. Tea Mosquito Bug
ii. Red Rust
iii. Grey Blight

The fourth category of classification was healthy tea leaves. The images were taken with a high-resolution smartphone camera and therefore, there were plenty of pixels of work with when reducing the resolution of images during model training (most deep learning models crop / scale images down to less that 300 by 300 pixels). A set of pre-decided guidelines were followed during the process of image collection:

1. As the first rule, the images taken for training purposes should be similar to those the farmers are likely to use for monitoring purposes.
2. Images were taken from varied zoom levels to ensure visibility of multiple individual leaves/ stems. Zoomed in images of single leaves that are affected by the pest/ disease infestation were taken.
3. The images of the tea leaves were taken in their natural arrangement rather than against a controlled background. The reason for leaving them in situ is so that the model can learn to identify the pests/disease with high accuracy when used in the field itself rather than being used necessarily in a controlled environment.
4. The images were free of any artifacts/ distortions like timestamps, etc. This is so that so that the model does not learn any unintended features.
5. The following metadata was to be recorded for each class of images so that after initial testing, the model can build on this information:

- Geographic Location (latitude/longitude or descriptive)
- Type of Crop
- Maturity of crop
- Type of pest or disease
- Description of stage of development of the infestation
- Zoom category
- Time of day
- Ambient weather (overcast, sunny, etc)

The images were internally classified according to the stages of infestation. However, for initial model training and testing, only the classification at the pest/disease level were focussed upon. The resulting image dataset consisted of approximately 500 images for each class taken at varying zoom levels, angles and sunlight intensity.

Examples of images at zoom levels 2 and 4 are shown in Figures 1 and 2 respectively.

**Fig. 1.**
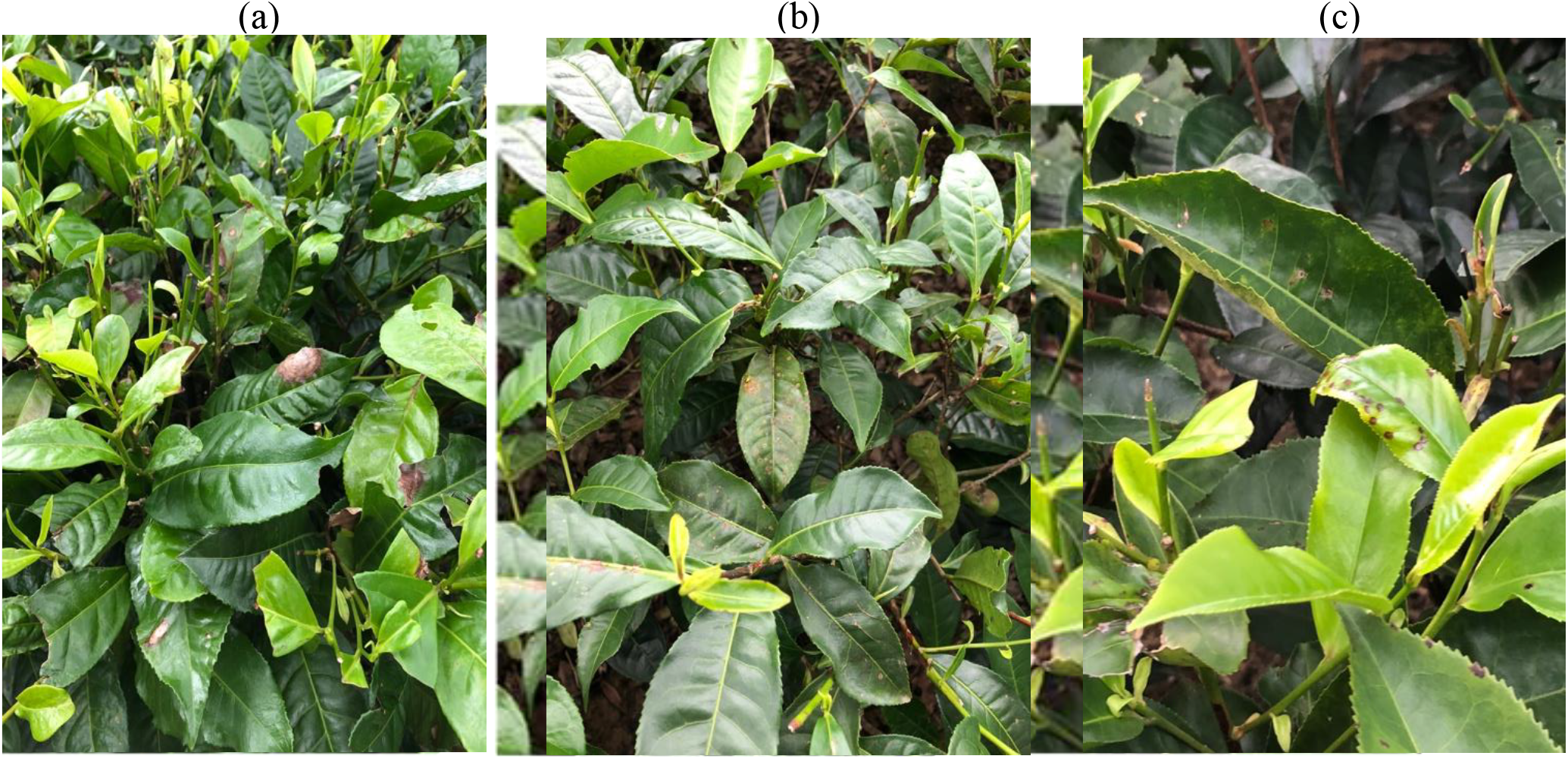
Examples of images at zoom level 2 showing (a) grey blight, (b) red rust, (c) tea mosquito bug

**Fig. 2.**
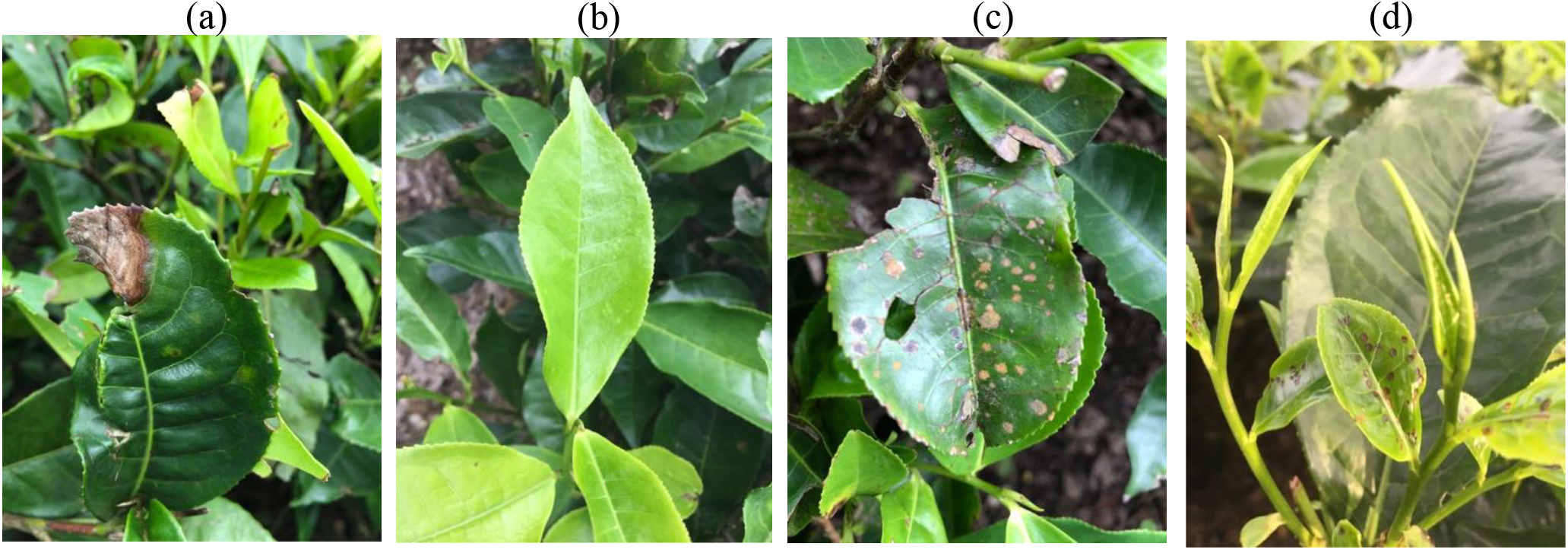
Examples of images at zoom level 4 showing (a) grey blight, (b) healthy, (c) red rust, (d) tea mosquito bug

For implementing the planned framework, a library called FastAi was used which is built on top of the deep learning package PyTorch. **PyTorch**, in common with other deep learning frameworks such as TensorFlow, provides what is called a model zoo. This is a set of models that have been developed for specific purpose and trained on large datasets so that they can be used with very little work provided that the task at hand is similar to that for which they have been developed. In the case of image recognition and classification there has been tremendous advances over the last six years since AlexNet (Krizhevsky *et al.* 2012). developed the first deep learning neural network to outperform traditional computer vision techniques. Such models are often trained upon a large corpus of over a million images, consisting of 1000 categories or more, called Imagenet (Russakovsky *et al.* 2015). With the weights learned from this training these models are then available in the model zoo and can be used to classify any image into one of the categories used in the training.

### Transfer Learning

Domain adaptation is a common requirement since often the data where labelled information is easily accessible and the data encountered in the application domain are different (Ruder, online). The model needs to be able to disentangle the general features and use that as an initialization to learn the domain specific features. Deep Learning models learn objects by learning features, and the different layers of the model recognise different types of features. Layers near the input learn low level features such as edges, gradients etc. Layers deeper in the model recognise how such low-level features come together to create objects which are specific to the images. What has been discovered over recent years is that these feature extraction layers can relatively easily be re-trained to recognise objects different from those they were originally trained upon, and the re-training needs far fewer images than the original training - typically in the order of hundreds rather than millions originally used. (Tan *et al*. 2018).

The process of taking a model that is pre-trained and adapting it in other domains is called transfer learning. The FastAi framework allows the user to very quickly and easily apply best practices in transfer learning. That is the basis of the work carried out here where a pretrained model is re-trained to classify the images in the dataset that contains the different types of pest/ disease symptoms in tea leaves. The pre-trained model used in this work was Resnet 34. There are many versions of Resnet and indeed many other alternatives such as ResNext, Inception, SqueezeNet, Densenet, MobileNet etc. Resnet was chosen since it provides a good balance between computational requirements in terms of memory, processing power and accuracy. The approach followed here could be repeated with any of the other models and would achieve still greater accuracy with some of the more complex models, however, the aim here was to establish the methodology first and Resnet34 provides a good platform to accomplish that.

### Resnet34: Architecture Overview

The winners of the ILSVRC 2015 image classification challenge introduced a new concept called a deep residual learning framework that made it possible to train hundreds of layers and still achieve compelling performance without worries of performance saturation (Bengio *et al*. 1994). Residual networks allow the training of deep networks by adding layers and adjusting the parameter values such that the additional layer does not change the output of the layer from which it receives input. This is known as identity mapping (Kaiming He *et al.* 2015; A. Ramcharan *et al.* 2017).

The training error of the overall module (with the additional identity layers) is the same as the error would be without the additional layers. Therefore, the additional identity mapping layer is used only for feature learning on top of the already available input. Since the additional layer is learning only the residual, the whole module is called a residual module. Resnet34, as used in implementation outlined in this paper, is a 34-layer deep residual learning framework and is illustrated in Figure 3.

**Fig. 3.**
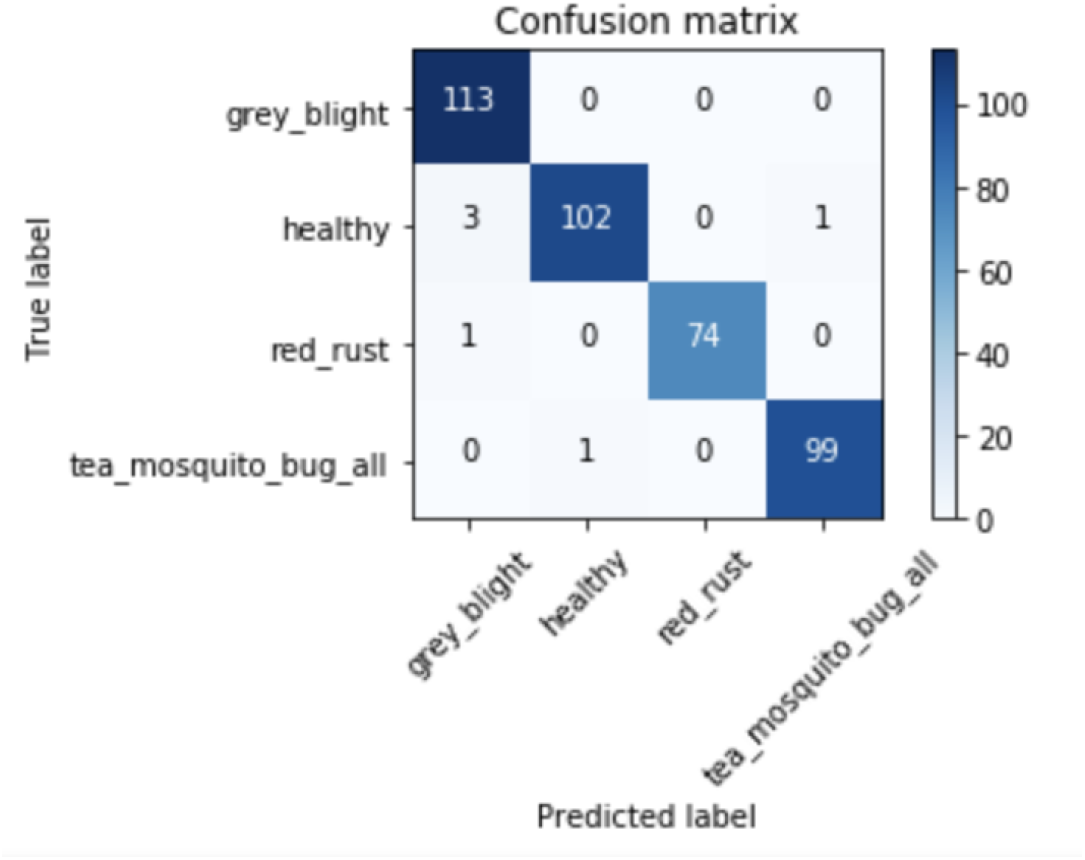
Confusion matrix for analysis

The architecture of a Convolutional Neural Network consists of 4 main operations which are the basic building blocks of every CNN (Goodfellow *et al.* 2016). These are:

1. **Convolution**: The primary purpose of convolution is to extract features from an input image. The convolution operation occurs between a filter (usually 3×3) and the pixels of the input image and an activation or feature map is produced. Different filter values extract different features (edges, gradients, etc) from the image.
2. **Non-Linearity (ReLU)**: Element wise operation that replaces all negative pixel values in the feature map by zero.
3. **Max Pooling**: A spatial neighbourhood (for example a 2 × 2 window) is defined and the largest element from the rectified feature map, within that window, is retained. This reduces the dimensionality of each feature map. Note that in many modern architectures the Max Pooling layer is being replaced at some or all locations by Global Average Pooling layer that effectively, instead of taking the maximum value in each area instead takes the average value.
4. **Fully Connected Layer**: The output of the convolutional and pooling layers are high level features of the input image. Fully connected layers at the end of the model use these features for classifying the input image into various classes based on the training dataset. The output layer uses a special layer based on a Softmax function so the sum of the output probabilities is unity.

The above CNN architecture is common to all models like Alexnet, Resnet, VGG etc (Krizhevsky *et al.* 2012; Kaiming He *et al.* 2015; Simonyan and Zisserman, 2014). The FastAi library groups the layers in the pre-trained model into 3 groups. Two groups are the convolutional layers (divided into two) and the third group is the fully connected layers part (J. Howard and R. Thomas, online).

### Convolutional Layers Feature Learning

The higher convolutional layers in the Resnet34 model capture generic features like edges, gradients, colour blobs while the deeper convolutional layers capture features that are more complex and specific to the dataset. The final fully connected layers capture information that is relevant for solving the specific classification task. Therefore, the approach while fine-tuning the weights in the pre-trained model would be to make significant changes in the weights of the deeper convolutional layers but smaller changes in the higher layers.

#### Cost Function

The cost function is a wrapper around the model that guides the calculation of changes in model parameters during model training. In this case the FastAI model automatically used the PyTorch Negative Log Likelihood (NLL Loss) function. This is commonly used for classification problems where there are more than two classes.

#### Gradient Descent

An optimization algorithm is used to minimize the loss function by iteratively moving in the direction of steepest descent as defined by the negative of the gradient. In the implementation outlined in this paper, gradient descent is used to update the parameters of the model to fit the tea dataset. Parameters refer to weights and bias values of each layer in neural networks. The goal is to adjust the parameters is such a way that the cost function is minimised. The way to do this is through calculation of the sensitivity (called the gradient) of each parameter upon the loss (effectively a measure of the error in the predictions). Given these gradients each parameter is adjusted to effect a reduction in the loss. The model is then run again and the process is repeated until a local minimum is reached and/ or the accuracy attained is adequate.

#### Learning Rate

The size of the adjustments made to the parameters in each step is called the learning rate. With a high learning rate, more ground can be covered each step, but there is a risk of overshooting the lowest point and the system becoming unstable. With a low learning rate, the cost function moves in the direction of negative gradient in a precise manner, however this approach is time consuming and it takes long to approach the local minimum (Krizhevsky *et* al. 2012; Bottou, 2012).

#### Batch Size and Epochs

The training set is divided into small subsets called mini-batches to update the parameters. However, to prevent any loss in accuracy and generality of the model, the model is trained on the whole dataset, divided into mini-batches, for several epochs. The batch size should be fixed in proportion to the image dataset i.e. small datasets should be assigned a smaller batch size (B. Kunwar, online).

## Re-training the Pre-trained model

### Initial Run without unfreezing layers

An initial run is done to check how well the pre-trained model can be generalized to the collected dataset and test how effective the dataset is for this purpose. It also helps in getting a rough estimate of the maximum accuracy that can be achieved with the dataset. The procedure is:

Step 1: Partition the dataset into train and validation sets.
Step 2: Centre crop of the image after scaling down to 300 pixels minimum for any side
Step 3: Use the FastAi library transformation features to apply random lighting, rotation and flip transforms and zoom levels. This augmentation of the dataset has a significant bearing on improving the accuracy of the model and also helps to reduce overfitting.
Step 4: The largest learning rate that can results in the fastest reduction of loss from a plot of loss vs. learning rate is selected (Bengio, 2012). In this implementation a value of 0.01 was chosen.

### Learning Rate Adjustment

As mentioned above, training with low learning rate is more reliable, but optimisation takes time because the steps taken towards the minimum of the loss function are small. Training with high learning rate may not converge as the weight changes are so large that it overshoots the optimiser overshoots the minimum and makes the loss worse. To overcome these issues, two features incorporated in the FastAi library are used:

1. **Learning Rate Annealing:** As the model get closer to the local minimum, the learning rate should be reduced. This allows taking of larger steps in the early runs, thereby saving computing resources, and allows more accurate solutions as a minimum is approached. The approach used is to assign a functional form to the learning rate. In FastAi’s implementation, the cosine form is used to maintain the learning rate high in the starting gradients and then drop quickly towards the minimum (Bengio, 2012). The learning rate is changed for every mini-batch, i.e. for every epoch.
2. **Stochastic Gradient Descent with Restarts (SGDR):** In some weight spaces, small changes in weights result is large changes in loss. The goal is to enable the model to find a weight space which is stable and accurate. This is done by a periodically imposing a large increase in the learning rate (‘restarts’) which will force the model to jump into a different part of the weight space if the current area is not resilient (G. Huang *et al.* 2017). The number of restarts is referred to as the number of cycles. The number of epochs between resetting the learning rate is set by the parameter cycle_len.

## Initial run Results and Insights

Using the learning rate finder provided by the FastAi library, the learning rate was fixed at 0.01 to carry out the initial run of the model according to the procedure above. The pre-trained model was run for 4 epochs. The accuracy at the end of the 4^th^ epoch was 89%. It was observed that this result was obtained without retraining any of the pre-trained features or any of the weights in the convolutional kernels, the only training was related to the final fully connected layers that feed into the classification output. This result was very encouraging as it implied that after retraining the convolutional weights of the pre-trained model with the tea dataset, significantly high levels of accuracy would be achieved.

## Unfreezing and Retraining

### Differential Learning Rate

Most pre-trained model architectures have many layers and specifying a learning rate for every single layer may be complicated for future fine-tuning purposes. FastAi provides the concept of ‘layer-groups’, which are layers in the same general part of the network are grouped together. The fully connected layers are one group and the convolutional layers are divided into two groups. The learning rates for each group is passed into the model as an array of three learning rates, one for each group (Howard, online).

The next step was to unfreeze all of the remaining layers of the model so that they could be retrained once an initial training of the new fully connected layers had been carried out. Before the unfrozen layers are trained, the appropriate learning rates have to be defined. As mentioned above, the earlier convolutional layers in the Resnet34 model capture generic features like edges, gradients, colour blobs while the upper convolutional layers capture features that are more complex and specific to the dataset. For the lower layers, the learning rate can be kept low, for the upper layers, the learning rate needs to be higher. The pre-trained model was retrained with the following learning rates: [lr/9, lr/3, lr] where lr or learning rate = 0.01. The model was run 14 epochs and an accuracy of 96% was obtained.

#### Fine tuning attempt 1

During image collection, it was observed that the disease Red Rust co-occurred with Grey Blight in many locations and some of the images. Examining the results of the analysis showed that a lot of red rust was getting misclassified as grey blight. To overcome this issue, the dataset was re-examined and images which contained both red rust and grey blight were cropped to leave either one or the other. After running the model on the edited dataset, the accuracy was found to have increased up to 97% and there were fewer incidences of misclassification due to confusion between red rust and grey blight.

#### Fine tuning attempt 2

Next, the above steps were repeated, but changing the learning rate for the layer groups to [5e-4, 5e-3, 0.02]. This process resulted in an increase of accuracy to ≈ 98.4%. The confusion matrix in Figure 3 shows a significant number of healthy labelled images being misclassified as grey blight and thus worthy of further analysis by implementing multi-label classification. Figure 4 shows examples of correctly classified images.

**Fig. 4.**
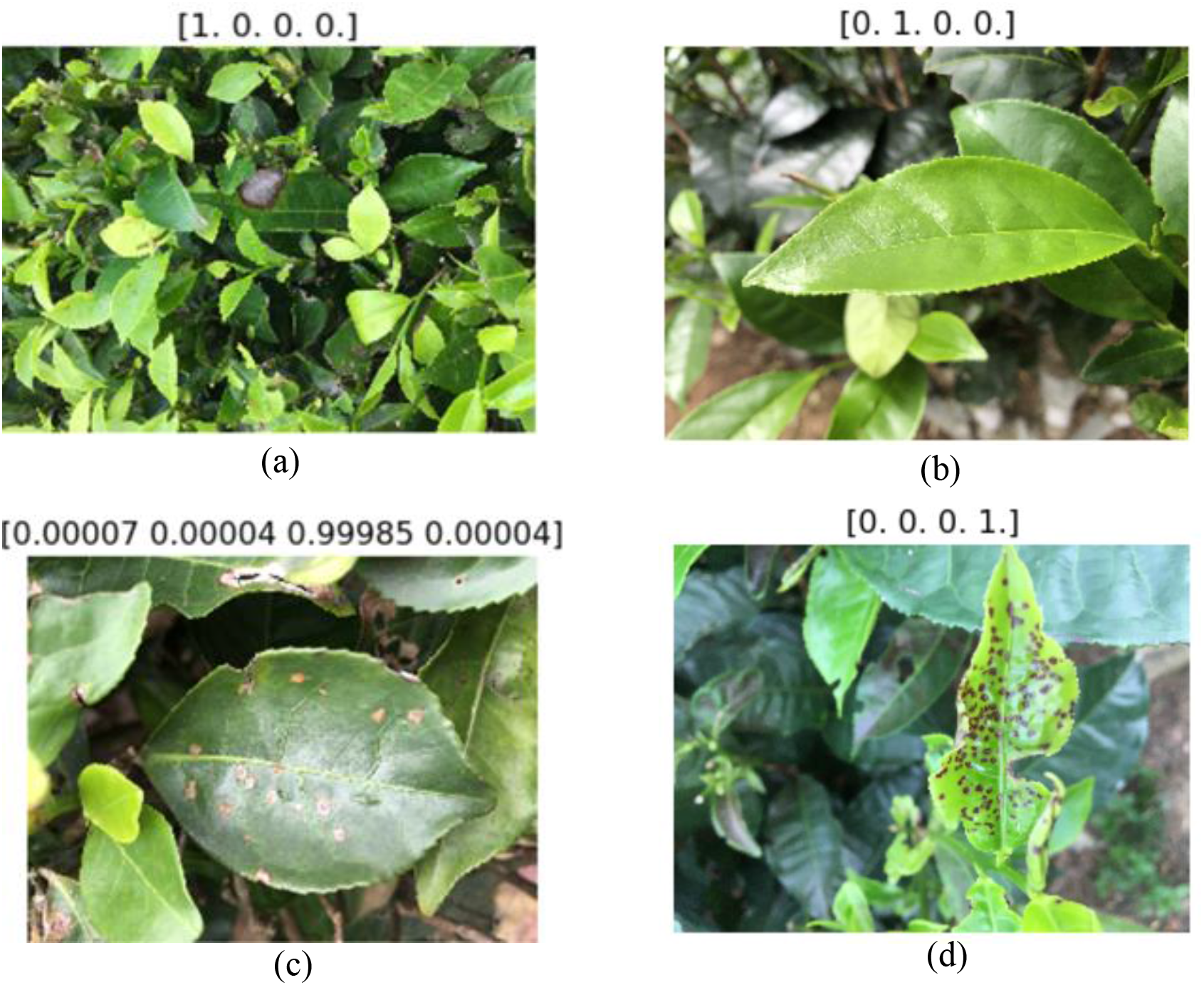
Examples of correctly classified images showing (a) grey blight, (b) healthy, (c) red rust, (d) tea mosquito bug

## 2.7 Incorrectly classified images

Figures 5 shows examples of incorrectly classified images. In Figure 5a this is actually grey blight, but it has been classified as red rust. One of the features of this image is that there is a lot of soil in the background.

**Fig. 5.**
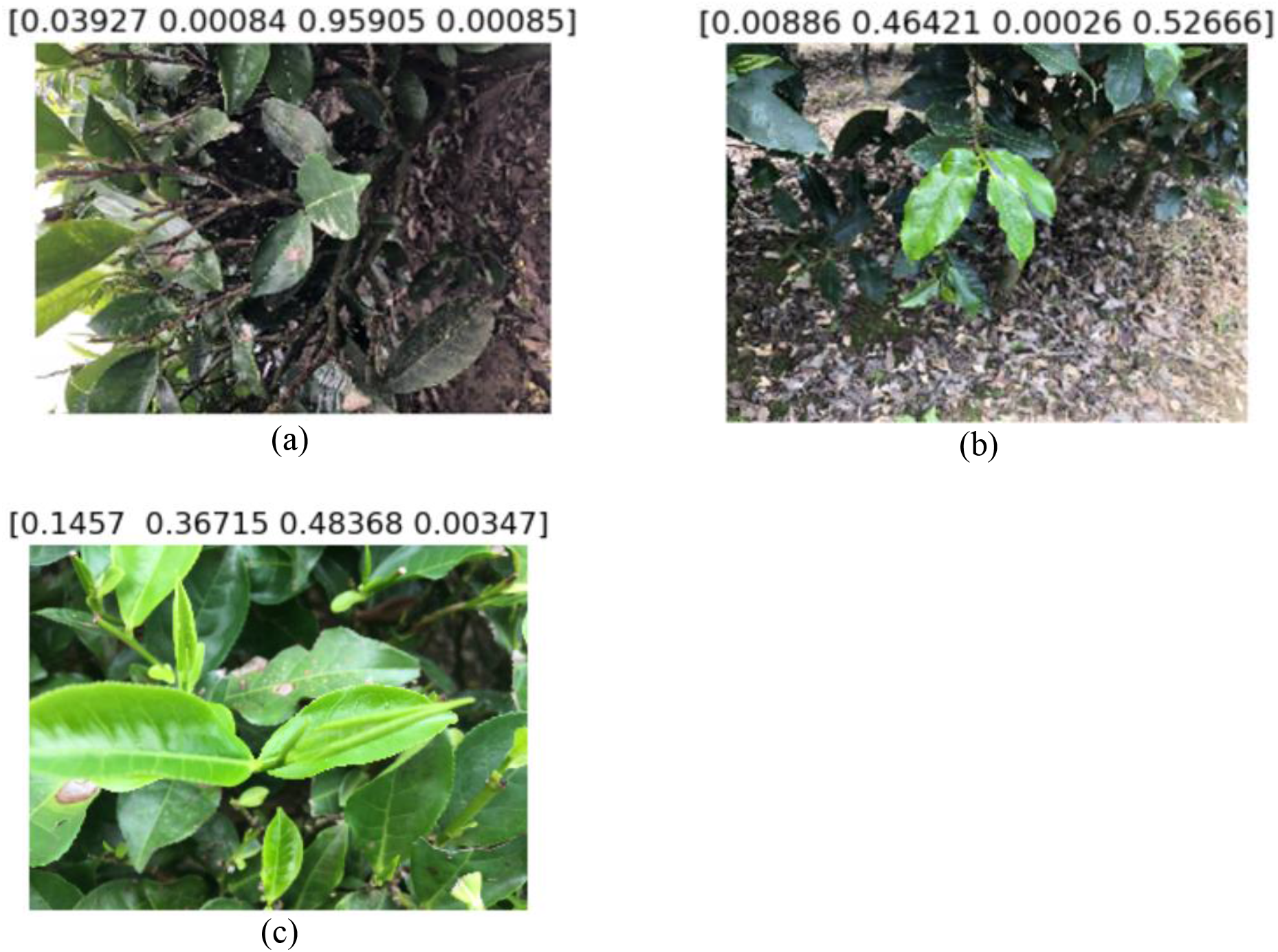
Examples of incorrectly classified images showing (a) grey blight, (b) healthy and (c) red rust and grey blight

After by looking at all of the misclassified images it was observed that that uncertainty or misclassification usually occurs in zoomed out images or in the ones where soil occupies a significant proportion of the image. It was also observed that zoomed in images also lead to correct classification with high accuracy (Noord and Postma, 2017). One reason for this is that the relatively low resolution of 300×300 pixels used in the classifier means that the fine detail of the disease is not clear in zoomed out images.

Another aspect of the dataset that was noticed while analysing the results from the above experiment was the images taken in the field contained leaves or set of leaves from different parts of the same plant and also the same set of leaves at multiple angles and zoom levels. This resulted in images containing leaves or set of leaves from the same plant in both training and validation sets due to the data splitting using a random process. This observation raised the possibility that the training process might be biased due to the presence of images of leaves from the same plant or plant group in both the training and validation sets. A well generalised model should have high prediction capabilities on the training set but also be able to generalise well over previously unseen data and this target is compromised by the effective leakage of the training data into the validation set (Reitermanov’a, 2010). A methodology was therefore developed to avoid this data leakage as outlined further.

From the above observations, it was hypothesised that in order to optimise the results there was a need to create a new dataset where the use of images that are zoomed out too far is avoided and to create a methodology for splitting out training and validation sets without the above-mentioned leakage.

The approach to overcoming these issues was:

1. As described above, the collected dataset consisted of multiple images of leaves that belonged to different parts of the same plant or group of plants at different angles and zoom levels. Therefore, the first logical step was to label the images according to the plant group to which they belong. This was done by assigning numeric codes to different plant groups (1, 2, 3, …, etc). The images belonging to the plant groups were assigned the respective plant group code. This ensured that when the dataset was split into train and validation, all the images from any group were in either the train or validation sets but not both.
2. Zoom levels were demarcated for each of the images again by assigning number codes for each level to maintain consistency. Images that are zoomed out have a lower numeric code, more zoomed in images have higher numeric codes. The next step involves modification of images which are zoomed out beyond a particular level and this helps in identifying the images that require modification.
3. Images with unrelated ambient content, e.g. soil occupying more than a reasonable proportion of the image, trees or fences in the background, etc required modification to remove the information that had the potential to influence the training process. In the images that were zoomed out to the point that more than one disease category was visible in the image, the image was cropped into two or more images so that only one disease or healthy category was visible in the image. The images produced by taking multiple crops of the original images were placed in the same plant group as the original image.
4. Finally, a master csv file was maintained to keep track of the changes being made to the dataset. The csv consisted of the following headers: Image File names, class, Plant Group, Remove code, Zoom Level, Modified. The Remove Code header refers to whether the original image has been divided and modified to the extent that the presence of the original image is irrelevant to the model training process. This csv was used as the medium to feed the images to the pre-trained model.
5. Once the images were ready using the abovementioned format, they were divided into training and validation sets using an algorithm that ensured any plant group was contained only in either the training or validation set

Once the new training and validation sets were produced, the same analysis process used for the original dataset was followed.

### Class Activation Maps

All the experiments undertaken above provided the insight that the images the model is exposed to in the field are complex, with myriad of attributes, distortions and ambient noise. This buttressed the need to validate the decision making of the network due to the complexity of the images that it was exposed to and understand the properties of the modules of the model network. This was done by visualising the weighted activations of the network for the images using class activation maps or CAM (Simonyan *et al*. 2013).

Since the model is trained using Resnet34 as the base model, the layers are not sequential but nested into modules followed by final pooling, dropout, batchnorm and fully connected layers ending with a log SoftMax (Ozbulak, Online; Erhan *et al*. 2009). The way visualization of the CAM was enabled was to change the model in the following ways. First, the model was cloned without the last 10 layers leaving an architecture that finished at the end of the last convolutional layer. A new convolution layer with 512 three by three filters but with only four outputs was added. Given the propagation of the 300×300 input image dimensions through the model this resulted in the layer output being four ten by ten arrays.

Following this layer, an adaptive average pooling layer was applied before finally flattening and applying to a log softmax. The model was then retrained by freezing all of the layers apart from the new ones, hence all of the original convolution layers remained exactly as they were. The final layers thus ended up with the 4 ten by ten outputs of the new convolution layer representing the feature maps or (CAM) of each of the final classes they were connected to.

These feature maps for each class are then overlaid on the input image in order to get the class activation maps. Note that it is important not to go back to the original image but to the image input to the model, since pre-processing will have resulted in cropping and other changes.

These class activation maps are helpful in assessing the importance of the image regions to the network activations and therefore, are very beneficial in identifying the discriminative regions and features of the leaves that activate the convolutional layers which facilitate the classification task (Selvaraju *et al*. 2016; Zhou *et al*. 2015). Mapping the activations for all classes are useful in understanding the reasons for misclassifications and if the model is getting confused between two classes due to similarity of features which is expected to happen between grey blight and red rust.

It should be noted that the exact classifications from this model might not be identical to that of the original model since the classification layers have been changed, however, the activations will be the same since all of the convolution layers are unchanged. In reality this modified classifier gives almost the same results in most cases.

## Results

### Correctly classified images of original dataset

In the following figures the numbers in brackets above each image show the predicted probability of the image content being of one of the four classes. A value of 1 indicated effective certainty that the content belongs to that class, a value of 0 suggests there is almost no probability that it belongs to that class. The class order is:

1. Grey blight
2. Healthy
3. Red rust
4. Tea mosquito bug

Figure 4 shows images that have been correctly classified with very high levels of certainty and Figure 5 shows one of the rare incorrectly classified images.

### Results of analysis of modified dataset

After training with the modified dataset an accuracy of 99.6% was achieved. This is better than the result of the previous analysis. This improvement is because the enhanced accuracy from the zoomed in images and removal of multiple classes in the same image acted to improve the accuracy and outweighed the reduced accuracy due to removal of data leakage. Overall, however, this model would be able to generalise much better on a potential test set. The confusion matrix for the modified dataset is shown in Figure 6 and it can be seen that only one incorrectly classified image resulted. Examples of correctly classified images are shown in Figure 7.

**Fig. 6.**
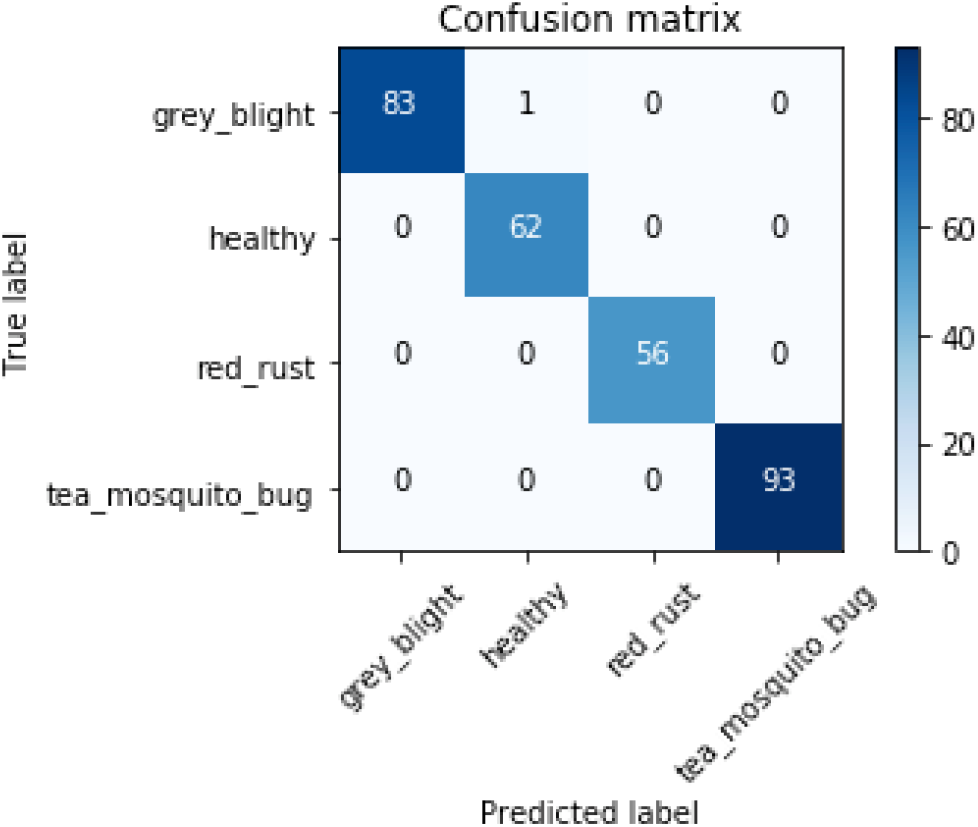
Confusion matrix for modified dataset

**Fig. 7.**
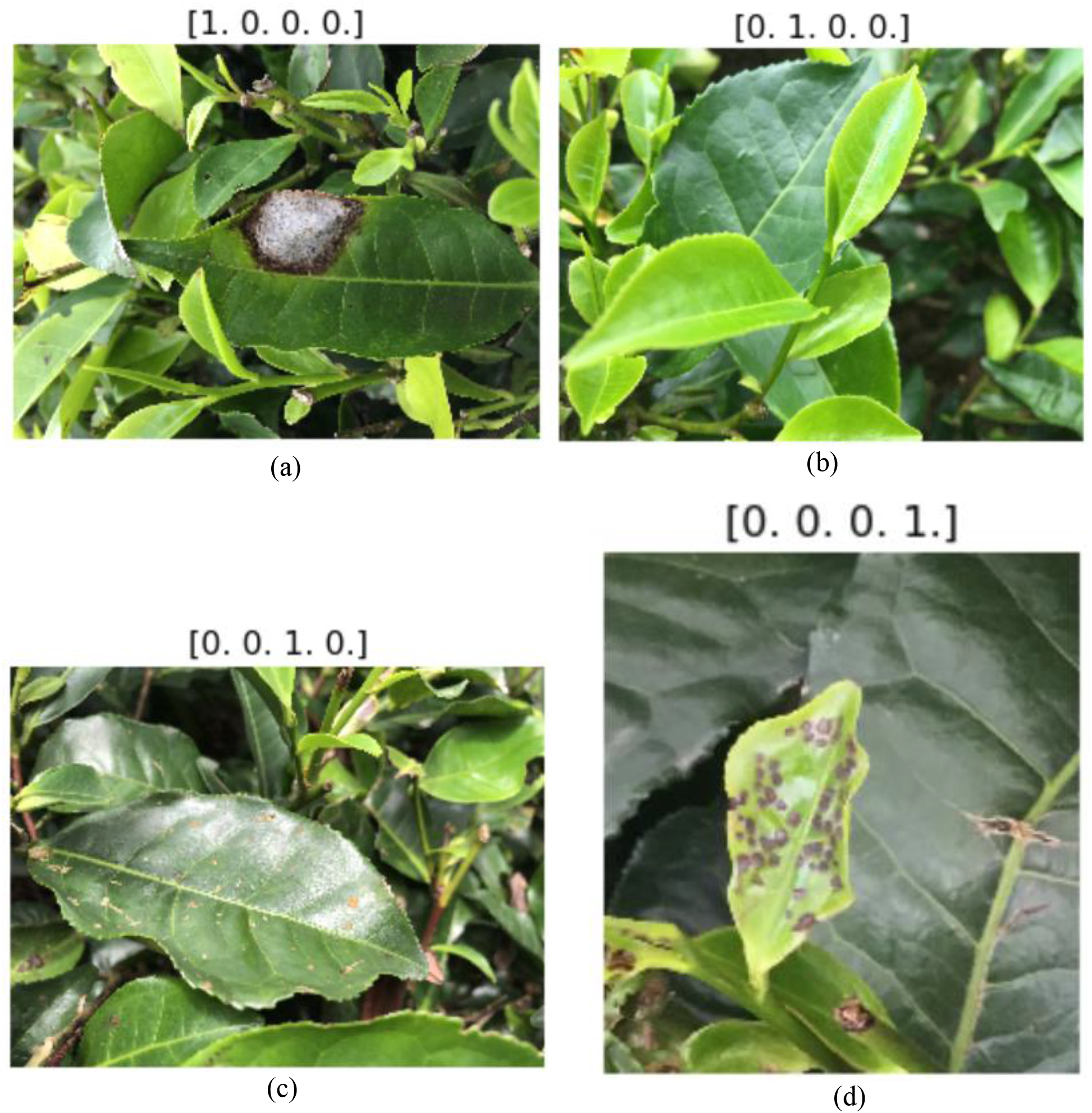
Examples of correctly classified images using the modified dataset (a) grey blight, (b) healthy, (c) red rust and (d) tea mosquito bug

Figure 8 shows the predictions for an image that has leaf damage. In previous models this had been incorrectly classified as grey blight. In fact, the leaves are either healthy or damaged but not diseased. This is correctly classified with the new model.

**Fig. 8.**
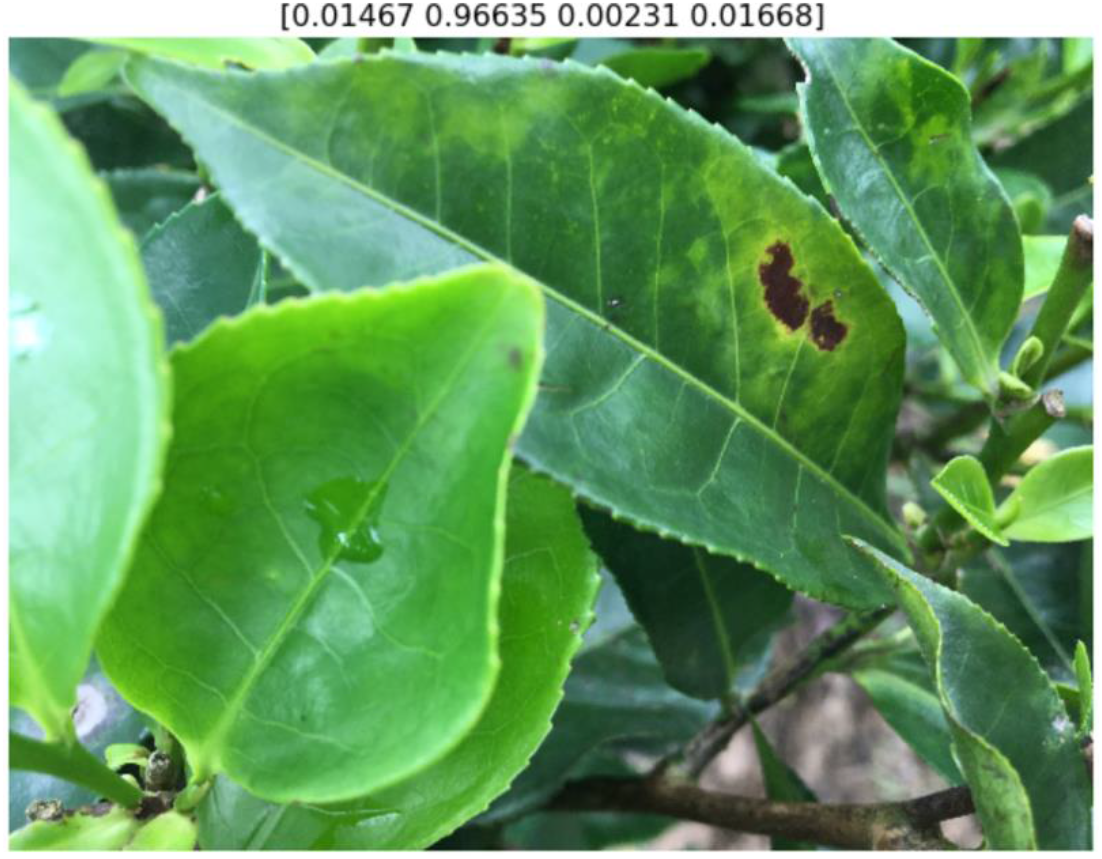
Correct classification of damaged and healthy leaves

### Class Activation Map Plot Analysis

In the following plots the title above each image refers to which of the CAM’s is being overlaid. In each case the order is according to decreasing final probability as per the last SoftMax are shown in the “Actual predictions” caption below the image. The non-linear nature of the classification process means that is not uncommon for the prediction probabilities to assume values of unity for one class and zero for other classes. Unless otherwise stated, to enhance understanding, the colour range of the CAM has been adjusted to cover the range from maximum to minimum over the displayed classes and the same scale used for all of the displayed images. Where there are very low activations for the other classes this often means that there is no significant colouring over the subsequent classes, which is why instead of showing the CAM plots for all four classes, in most instances, the plots for the two classes with highest prediction probabilities have been displayed.

Figure 9 shows prediction from leaves that have grey blight. The predictions show that the model has correctly identified the above image as grey blight and with a high degree of accuracy. However, the CAM for grey blight shows that apart from the prominent grey blight infected part of the leaf, the network is picking up a little bit of the leaf in the forefront (right) which is only damaged and uninfected, but the network could be confusing the damaged leaf to be symptomatic of grey blight because of the similarity in physical attributes.

**Fig. 9.**
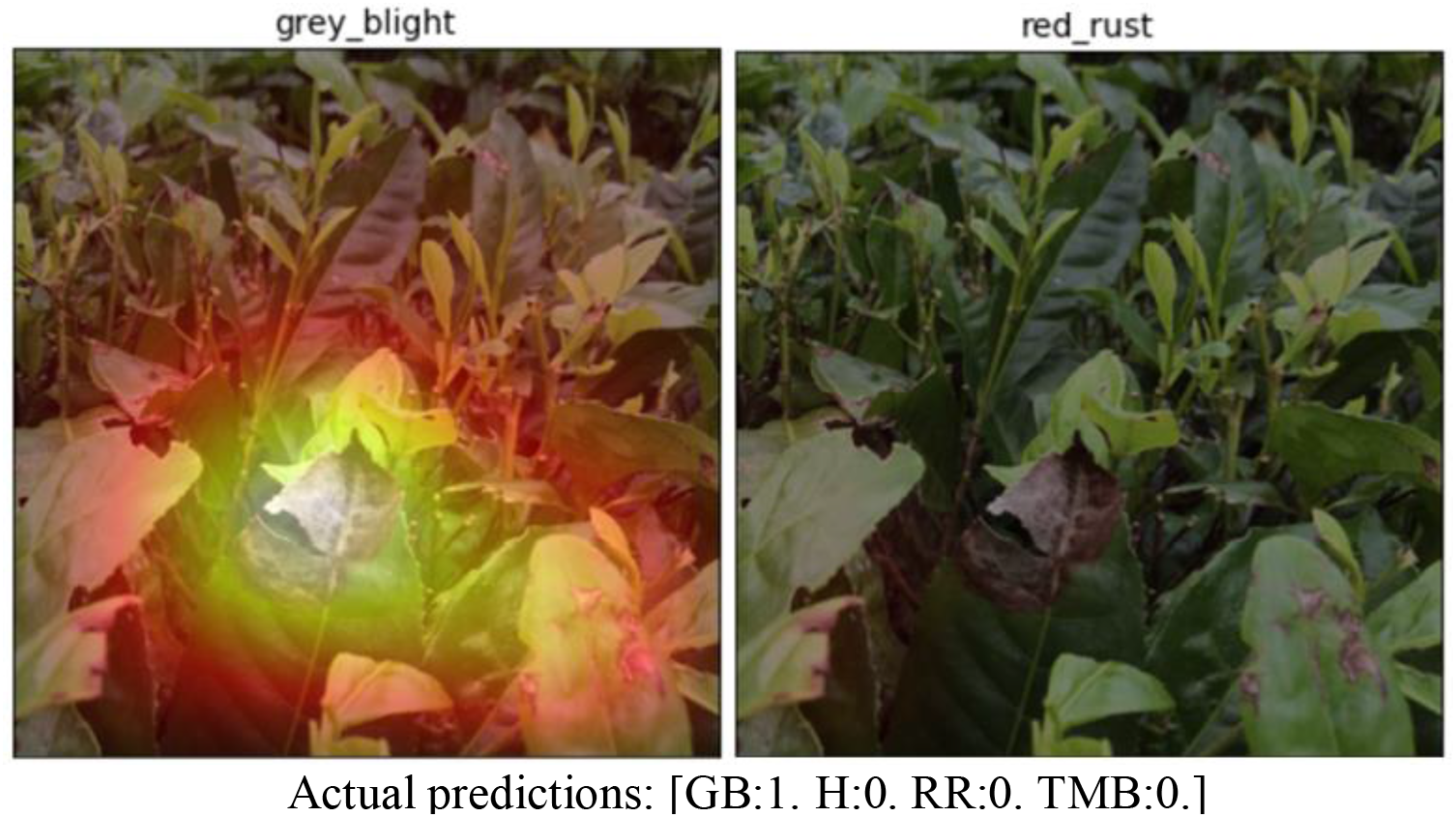
CAM for image IMG_2848

Figure 10 and 11 examine classification of the same leaves that are classified differently at different levels of zoom. Note that in this case the colour map scale was modified to focus upon areas with the highest activation levels since the default levels made most of the images a mid-intensity orange. Figure 10 is zoomed out significantly more than Figure 11. Figure 10 is the most worrying of all in that it is a clear misclassification. The actual disease here is grey blight, however the class with the highest prediction probability is red rust, despite this having relatively low levels of activation, although what there is cover a wide area hence giving a higher value from the global average pooling. In this case max pooling would give a different result, however, the result would still be a miss-classification since the class with the second highest probability is tea mosquito bug. This is probably because the two dark holes in the leaf could be mistaken for tea mosquito bug. It is noticeable that in the healthy image the good leaves on the right-hand side of the image are recognised. In the grey blight image, it can be seen that the larger area of grey blight is picked up correctly, but the activations are quite low suggesting the model is uncertain. Figure 11 shows the same leaf but enlarged and from an angle where the large grey blight is not partially obscured by the leaf in front. In this case the classification is accurate, and both areas of grey blight are identified correctly. This demonstrates that predictions should be based upon ensemble models based upon several images taken with variable viewpoint and magnifications, rather than just one image. This is a very important finding and conclusion.

**Fig. 10.**
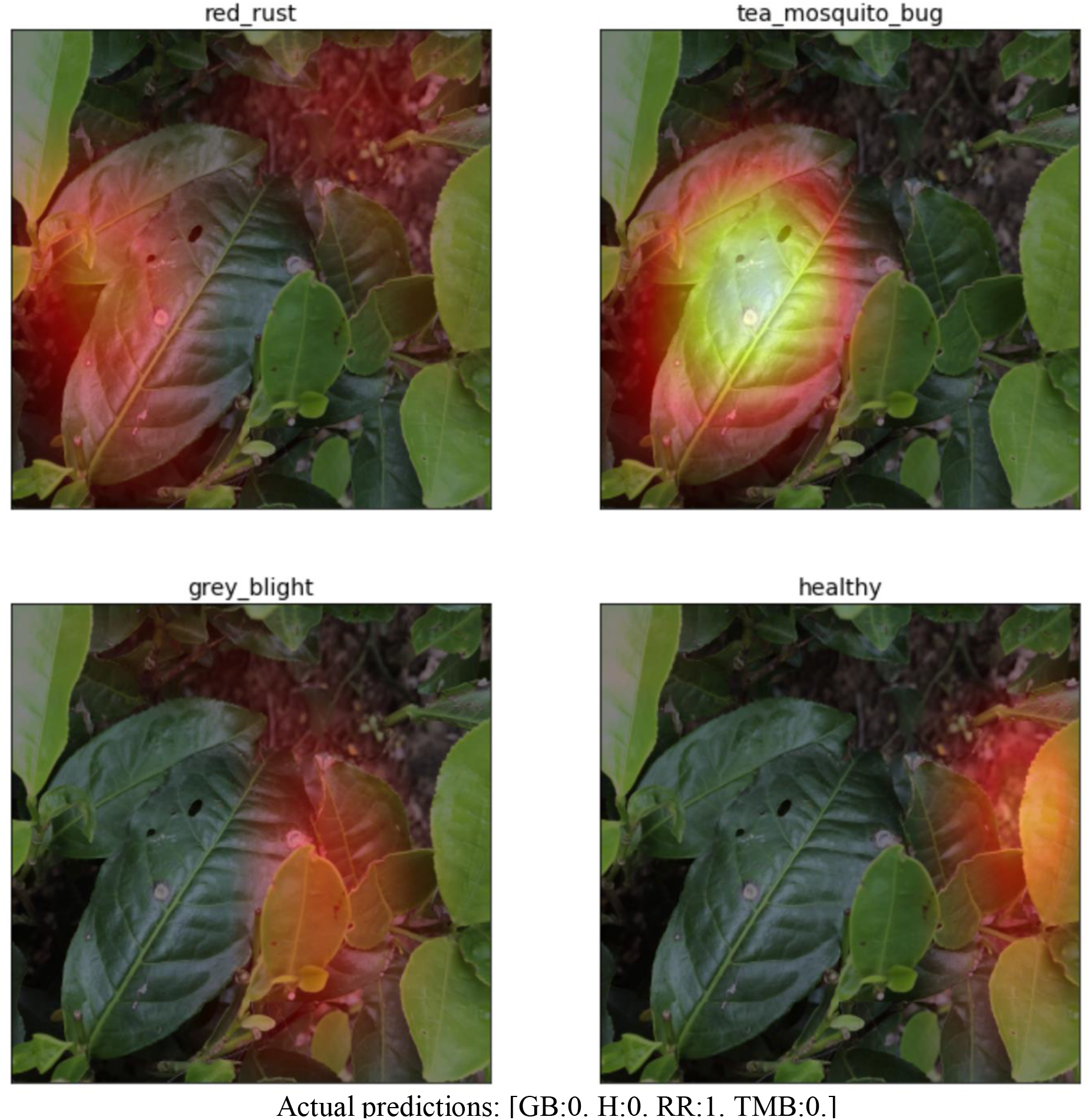
CAM for image IMG_2980 showing miss-classification

**Fig. 11.**
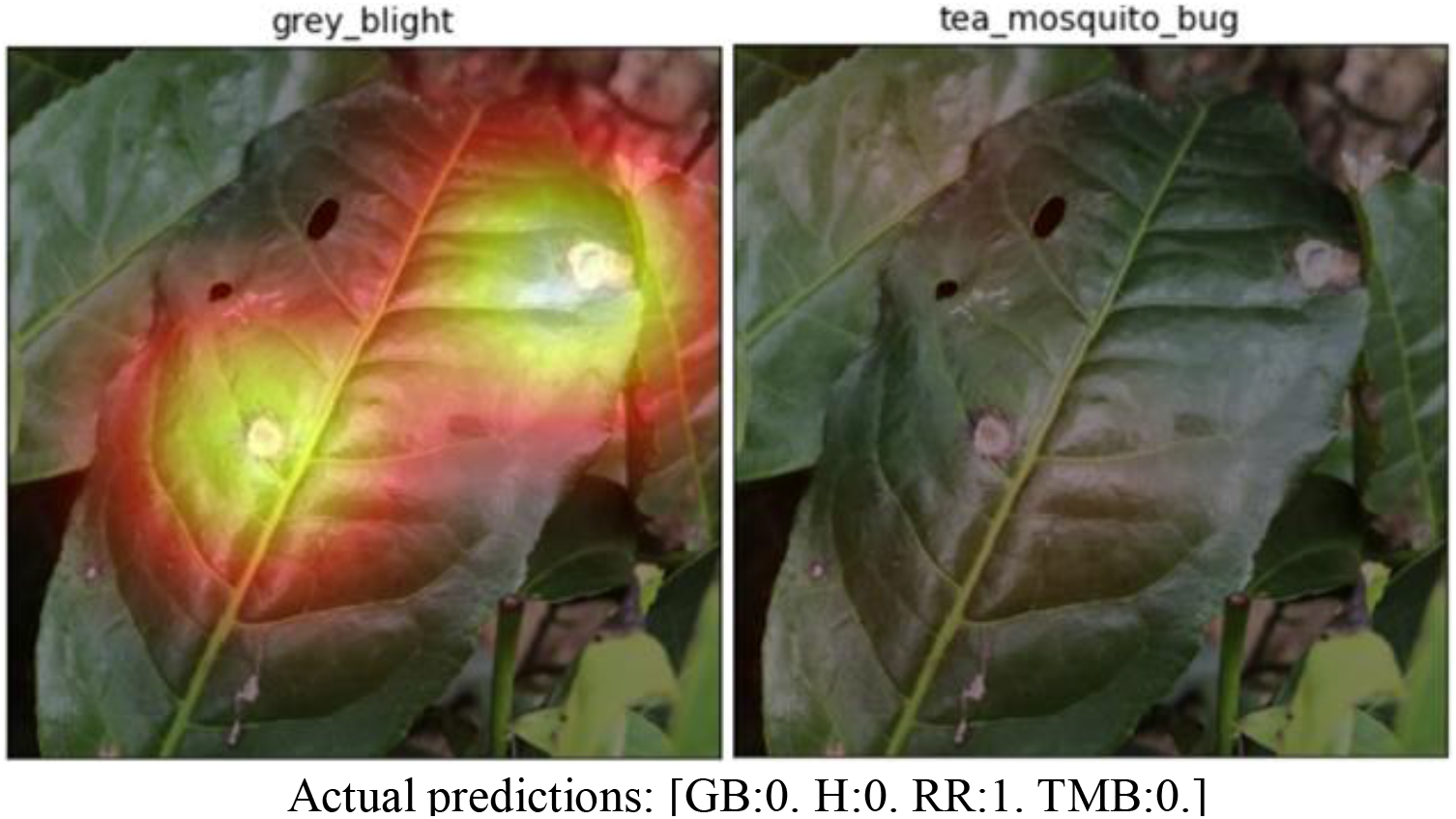
CAM for image IMG_2981 showing correct classification when the image is more zoomed in

Figure 12 shows an image where the centre leaf is correctly classified as grey blight infected. Interestingly, looking at the CAM for tea mosquito bug, the network does localise some features in the leaf on the right as indicative of tea mosquito bug. To understand this, the original image was analysed to find that the leaf simply has some black spotty coloration which can look in some respect similar to tea mosquito bug but is not. This confusion of features is an added complexity of working on an image dataset collected in the field. The CAM visualisation goes a long way in providing these insights.

**Fig. 12.**
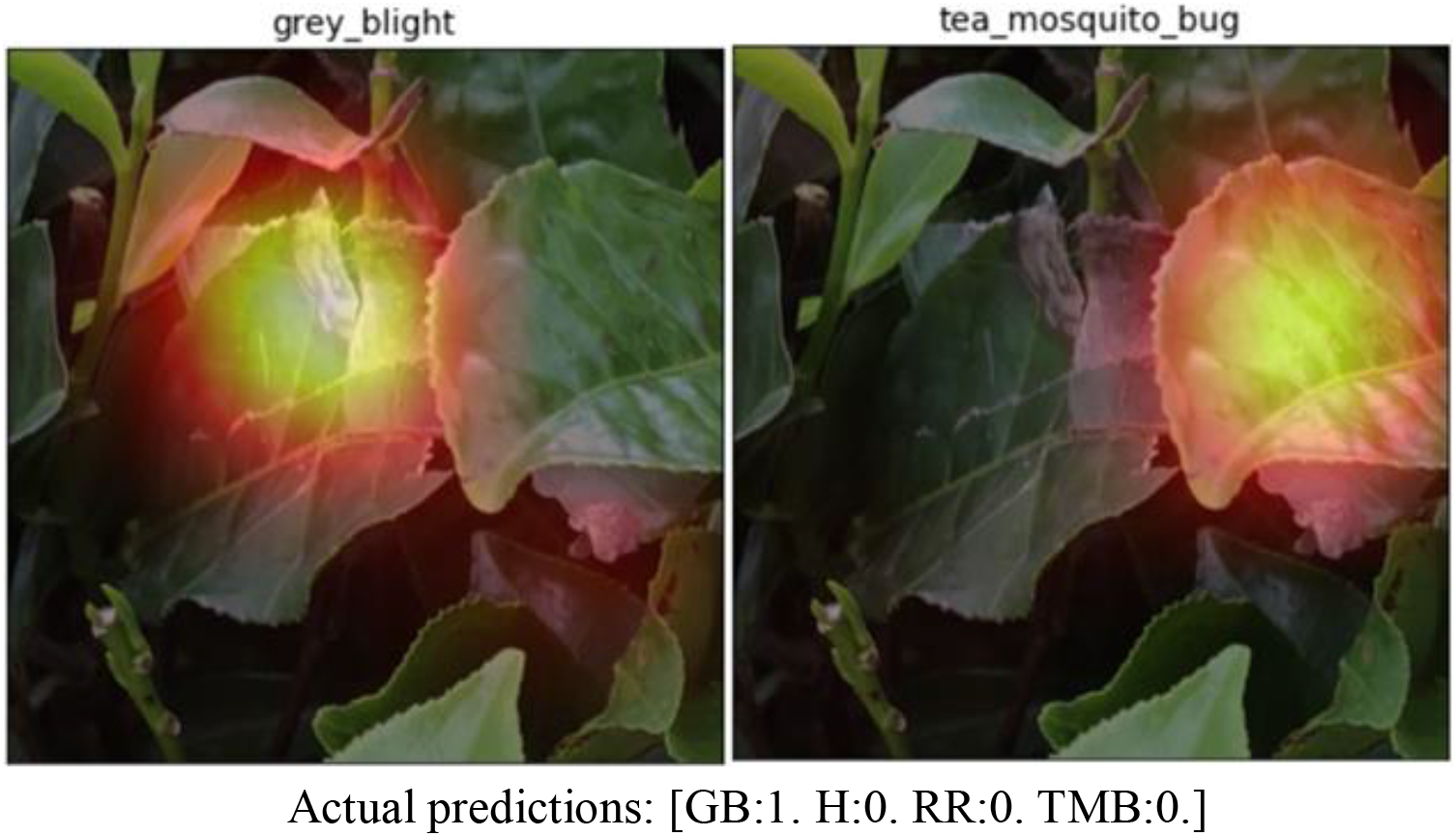
CAM for image IMG_2983 showing correct localization of grey blight. It is noticeable that the light black dots on the leaf on the right are seen as potential tea mosquito bug

Figure 13 shows an image that is correctly classified as grey blight, however, there are several leaves in this zoomed out image that are infected with grey blight but the CAM localisation shows that the convolutional network is being activated primarily by the large grey blight features present in the leaf in the foreground and is ignoring the others that are present in background. Therefore, in the field, the model will prioritise features of the leaves in the foreground and in focus. It is likely that the model gains in confidence that larger areas of grey blight are indeed grey blight, but lower confidence in smaller areas that could be confused with tea mosquito bug or simply damaged leaves in otherwise healthy leaves.

**Fig. 13.**
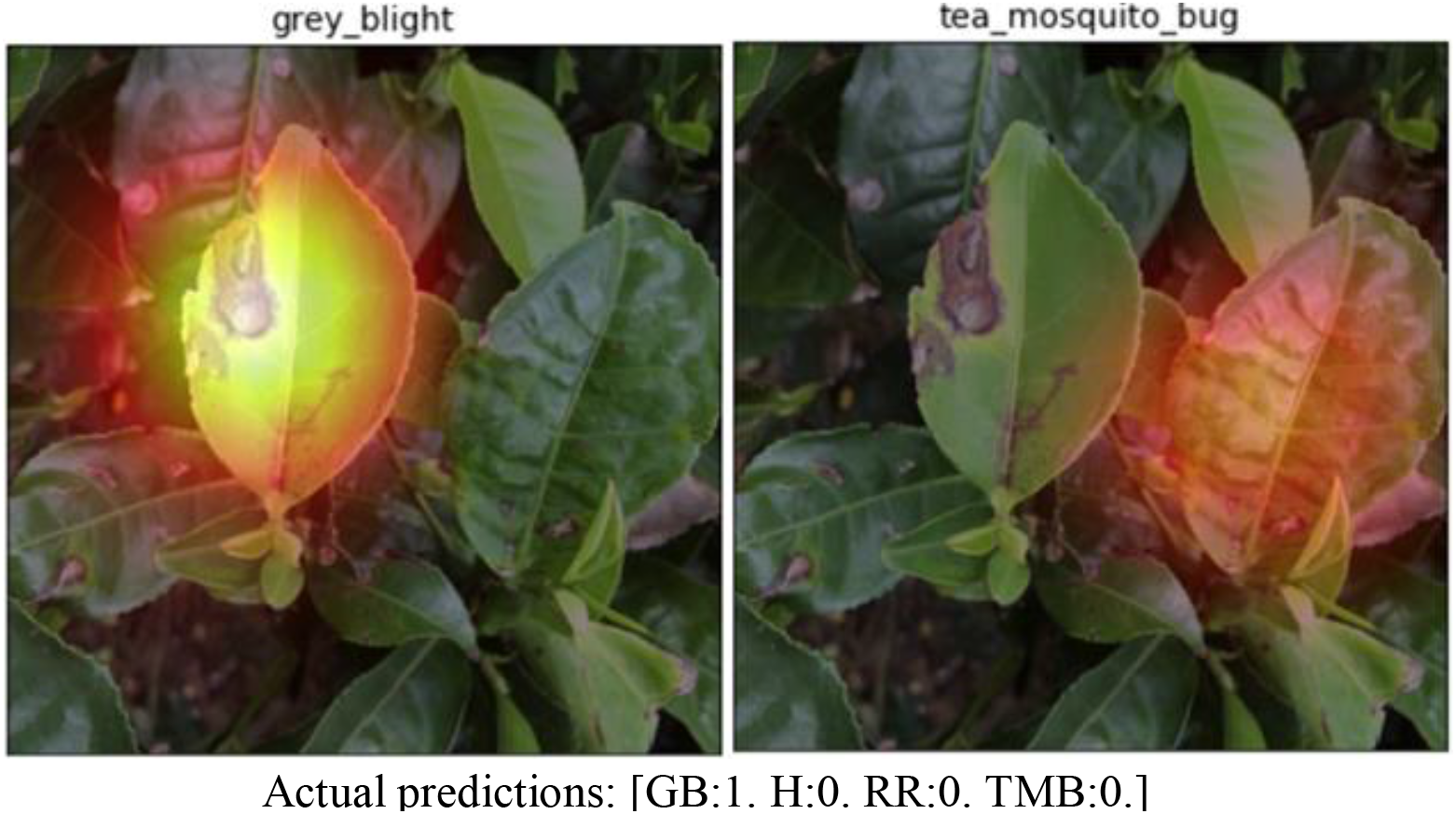
CAM of IMG_2984 showing detection biased to largest regions

Figure 14 shows an image where there are two leaves that are symptomatic of red rust. The localisation of the CAM shows that the network is successfully picking up features on both the leaves, even those features that are not very pronounced (red rust in early stages).

**Fig. 14.**
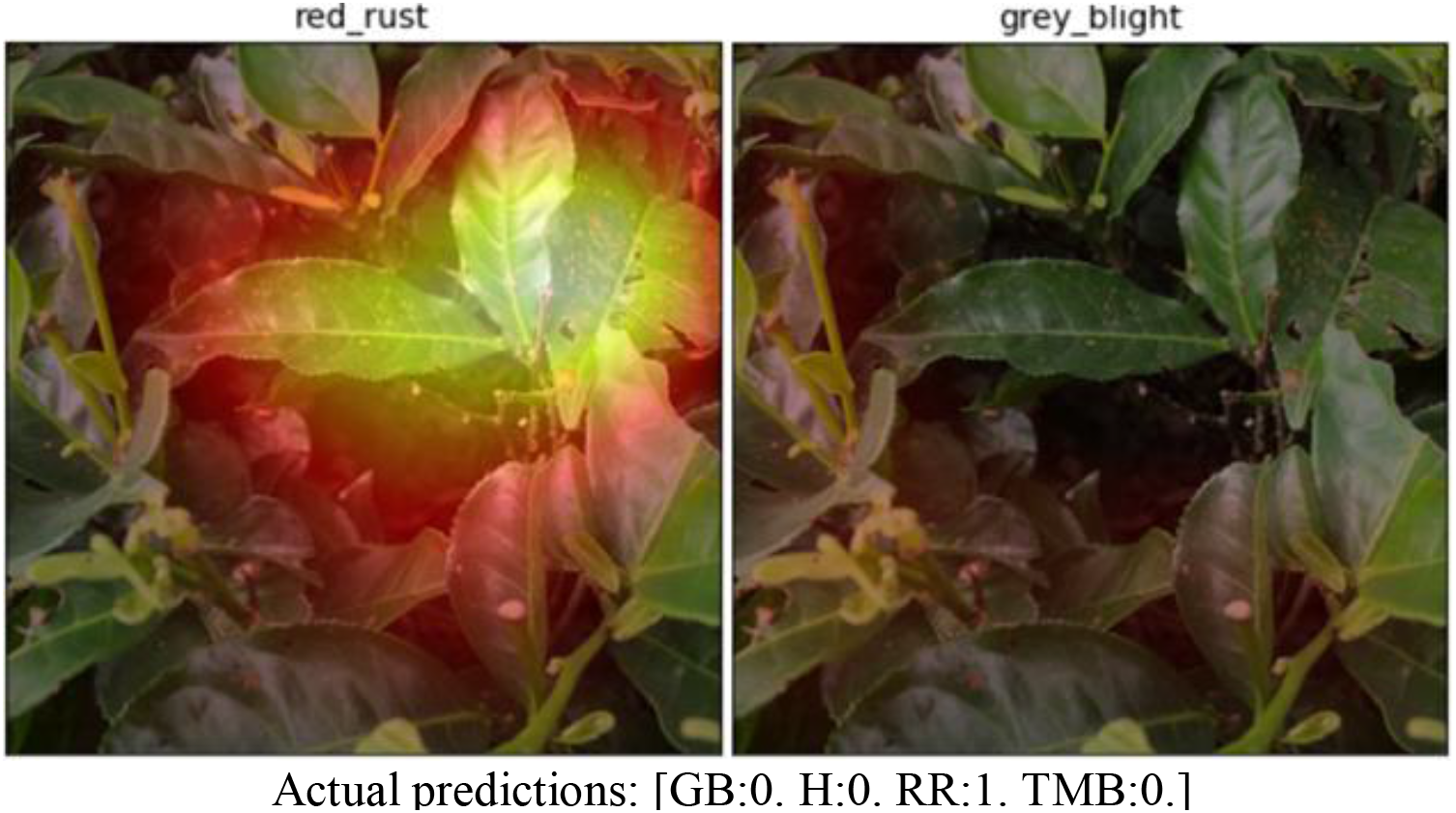
CAM of IMG_3217 showing correct identification of Red Rust on several leaves

Figure 15 shows CAM activations for one of the images that was found to have reduced confidence in the predicted class. In fact, it is correctly classified as red rust but with relatively low accuracy. The CAM visualisations show the reduced confidence is not due to a problem with the identification of the red rust, but instead due to the fact that all of the other classes are also detected in the image. Since the model is being forced to choose only one class the presence of a host of other features reduces the confidence level. This result is encouraging and lends support to the viability of developing the model into a multi-class prediction tool in the future.

**Fig. 15.**
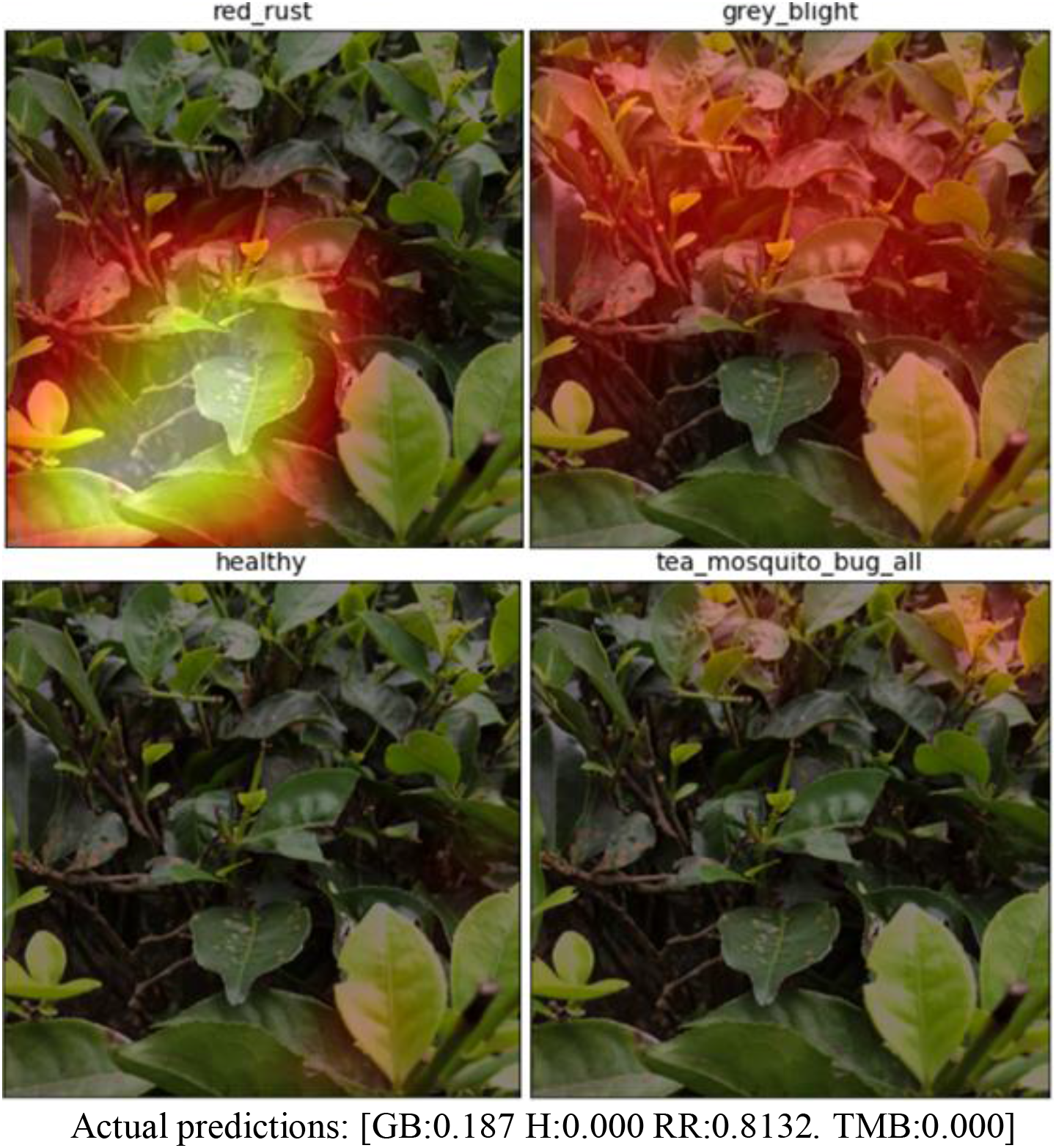
CAM of IMG_3415 showing how the presence of multiple classes can reduce predictive confidence

The image shown in Figure 16 has a lot of soil showing in the background. There was some concern that this would be detrimental to the analysis, however, it can be seen that the network is able to localise the features of the red rust and ignore the noise added by the soil.

**Fig. 16.**
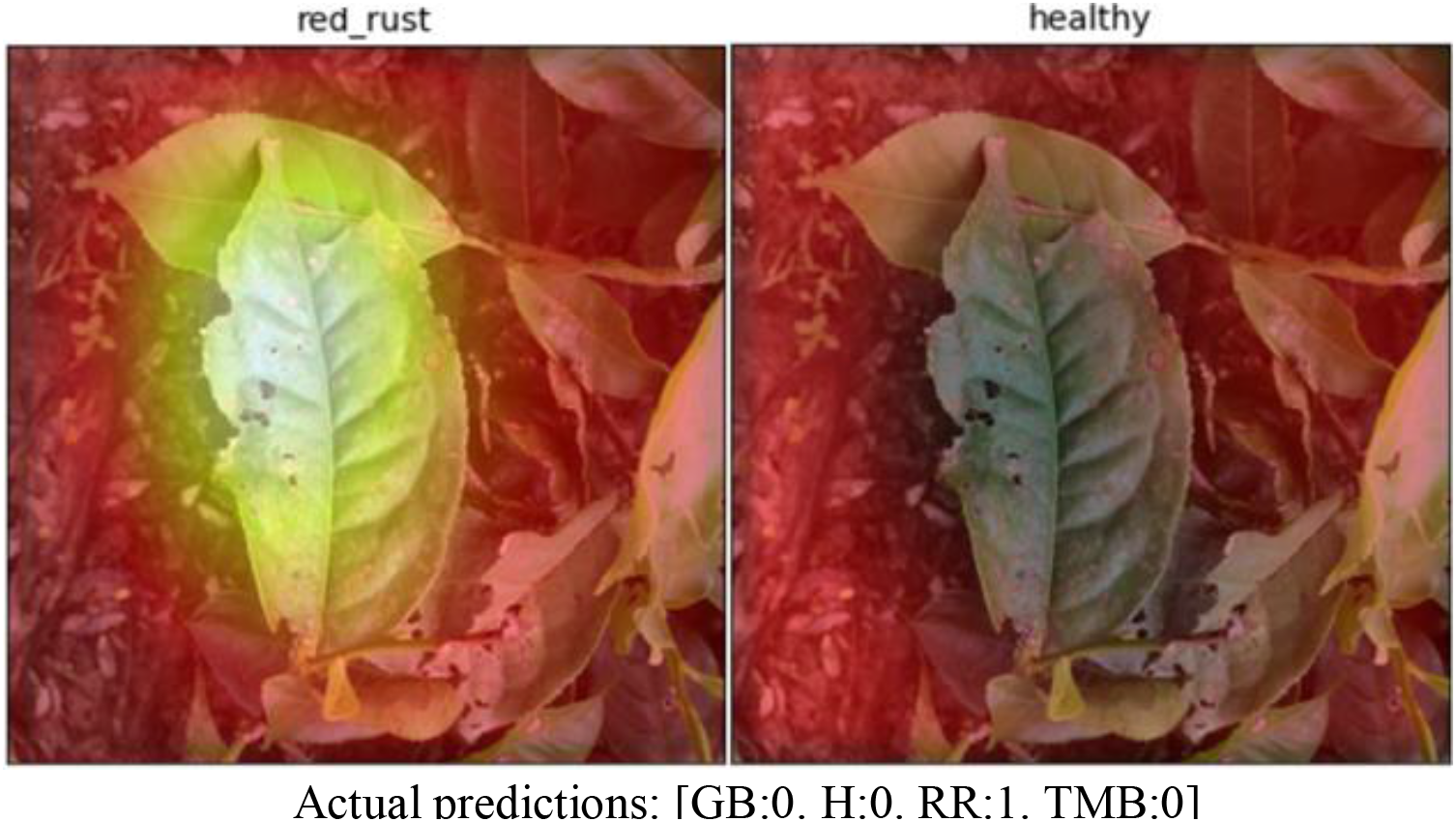
CAM activations for IMG_3430 showing that soil in the background is not detrimental

Figure 17 shows the convolutional network is able to sufficiently ignore the noise from the object in the background of the image and focus on the features of the leaf infested with tea mosquito bug. However, CAM visualisation does show a small amount of the background object being picked up by the convolutional network which shows that the presence of such noise impacts the way the convolutional layers recognise the features.

**Fig. 17.**
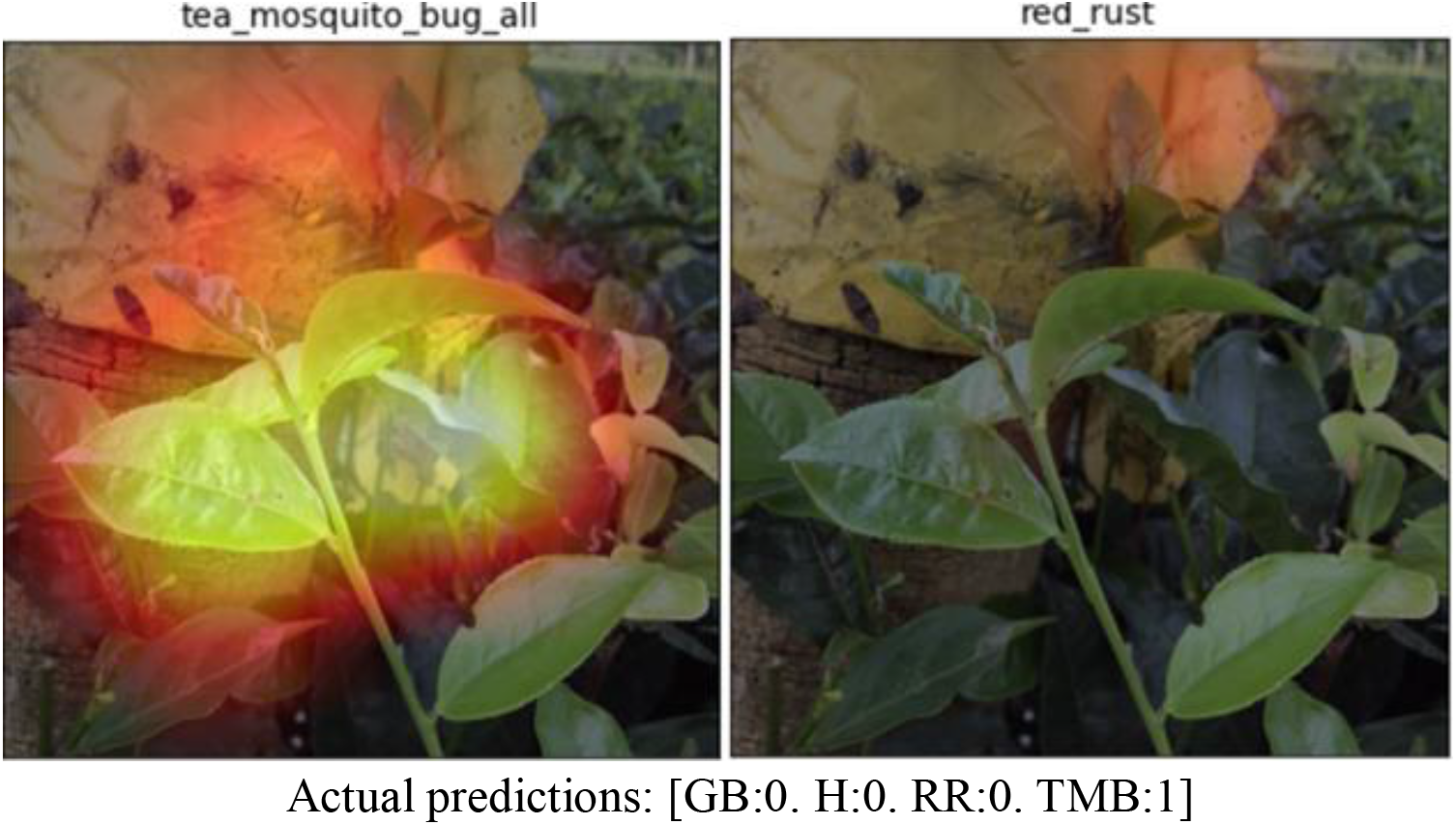
CAM for IMG_2159 showing detection of tea mosquito bug despite the presence of material in the background

## Discussion

This work was intended to investigate the feasibility of using deep learning image recognition tools for diagnosis of plant disease from smartphone camera images. The work has three stages: the original dataset; the dataset with groups; and the class activation mapping investigation.

### Original dataset

The model performed suitably well after it was trained on the original dataset and gave an accuracy of 98.4 % post fine-tuning steps such as learning rate adjustments for each group of layers. Despite this high accuracy, analysis of the confusion matrix showed a number of misclassifications between healthy and infected leaves. The reasons for this were believed to be three-fold, these were the presence of multiple diseases in the same image, the model focussing on leaves other than the target infected leaves and also that some of the images were zoomed out too far. There were also concerns about leakage between the training and validation sets. The high overall level of accuracy of this work gives confidence that the approach was viable and led to the methodology for development of the modified dataset.

### The Modified Dataset

The original dataset was modified by reducing the zoom level of highly zoomed out images and assigning images to plant group where there was no leakage of leaves from one group to another. The training and validation sets were created based on these groups. It was anticipated that certain biases might have crept into the model due to an imbalanced partition of the data into training and validation sets and therefore the dataset partitioned according to groups might face some loss of accuracy during validation. However, the accuracy after unfreezing and training with differential learning rates was 98.3% and after using an optimal learning rate of 0.001, became steady at 98.6%. The validation loss showed a steady decline, being greater than the training loss only by a small margin ≈ 0.02 so there was no fear of overfitting. It became germane to the discussion of training a model to be suitable for field use to understand where any remaining errors were coming from and to see what factors could reduce confidence even if they did not result in a mis-classification. This led to the work with CAM visualization.

### CAM Visualisation

The Class Activation Mapping visualisations helped identify the regions of the image that were being focussed on by the layers of the classifier and also enabled prediction of the way in which the model, when loaded on a system like a smartphone, would behave in the field. As evident from the results, the model identifies the symptomatic parts of the leaf in the case of an infection and is able to carry out the single class recognition task to a significantly low zoom level. From the highly zoomed images, the model is able to learn detailed features of the leaf and performs classification with very high accuracy. However, for images that are significantly zoomed out, the CAM visualisation shows that the model prioritises the leaves that are in the foreground. The visualisations also indicated that it is possible for the CNN to get confused when multiple leaves are present in the foreground causing the accuracies of the predictions to drop. The model is currently not optimised to differentiate between physically damaged healthy leaves and diseased leaves. In an image which high level of background noise, CAM shows that when the proportion of noise (example: soil) is below a certain value, the classifier successfully ignores the distraction and focusses on the learned features. However, when the proportion of the noise in the image increases (example: a black pepper tree along with significant part of the landscape), the prediction accuracies tend to drop. This is an idea that needs to be kept in mind when applying this classifier on the field level because the images that the classifier will encounter will contain multiple leaves and most likely have more than one diseased leaf in the same frame surrounded by many healthy leaves with high noise levels. From the CAM visualisations it can be estimated that the model will perform well in the field, however a multi class model would be more practical for a field level application.

The CAM visualisation also informs the image collection process. This entire project impresses the fact that working with leaves and their images in the field is a relatively complex process in comparison to training a model on a leaf dataset in a controlled set up and a noise free environment. As seen in Figure 10 a model trained on such a controlled dataset might work very well on paper but may fail on the complex and noise intense conditions of the field. Analysing and deciphering model behaviour based on CAM visualisations form a very important part of incorporating this model in an application that can be tested on the field level with reasonable results.

## Conclusions

Based upon the work to date it is considered that the approach adopted can be developed as a way to help farmers diagnose plant disease. Important learning points for the next phase are:

1. Analysis should be made using ensemble analysis of multiple images of the same leaves
2. The level of zoom should be such that features can be clearly recognised in a scaled down 300×300 pixel image, ideally with one of the diseased leaved occupying a large proportion of the image
3. Once the convolution layers have been trained it would be sensible to look to enable multi-class classification, which should be a relatively easy change to make and would allow the model to cope with features it commonly encounters
4. Although not mentioned here in the future region-based models such as Unet [ref] might be necessary to enable quantification of the stage of the disease (based upon for example size and number of disease areas). This should be possible but will require development of a training dataset with bounding boxes added as well as disease classification.
5. Alternative base models such as ResNext could provide greater robustness of classification albeit at the cost of increased training and application resources.
6. The use of Class Activation Maps to validate and understand what is being recognised by the convolutional layers is a very valuable tool.

## Supporting information

Title Page

## Acknowledgements

This research was supported by the Group Technology and Innovation Office (Tata Sons), Rallis India Limited and Tata Global Beverages. We are thankful to Vidya Hegde and Dr. Mallikarjunappa (Rallis India Limited) for providing assistance that was helpful in guiding the research. We express our appreciation for Kailyanjeet Borah (Tata Global Beverages) for providing the necessary support that expedited the process of data collection and expert knowledge about the tea crop and various pests examined in this work.

We are grateful to our colleagues Dr. Gopichand Katragadda, Kamesh Gupta, Piyush Mishra, Aditi Olemann and Arati Nijhawan from the Group Technology and Innovation office for providing wisdom and support that was instrumental in taking this research to its fruition.

## Notes

#### Summary of Updates

Formatting changes - removing line numbers in manuscript pdf.

