## Supplementary material for "The development of methodology and techniques for crop disease identification": Title Page

N. S. Tiwari^1^, J. W. Richmond^2^

^1^Boston University and Tata Group Technology and Innovation Office. Current: View, Inc

^2^Tata Group Technology and Innovation Office
